# Neuraminidase 3 acts in a rapid translation-based positive feedback loop to activate TGF-β1

**DOI:** 10.1101/2025.10.16.682941

**Authors:** Sumeen Kaur Gill, Richard H. Gomer

## Abstract

Fibrosis appears to be an out-of-control wound healing response that drives a progressive formation of scar tissue in an organ. A key profibrotic cytokine, transforming growth factor beta-1 (TGF-β1), upregulates levels of the fibrosis-driving sialidase, neuraminidase 3 (NEU3), and NEU3 can activate latent TGF-β1 to release active TGF-β1 from the sequestering LAP peptide. In the mouse bleomycin model of pulmonary fibrosis, NEU3 is both necessary and sufficient for fibrosis. In this report, we find that NEU3 levels increase both intracellularly and extracellularly in cultures of human lung fibroblasts within 5 minutes of TGF-β1 exposure. This effect is driven by an increase in translation and is independent of new transcription, supporting a model where TGF-β1 causes a pool of weakly translated *NEU3* mRNA to increase translation. The RNA helicase DEAD-box helicase 3 (DDX3) mediates NEU3 translation. TGF-β1 induces dephosphorylation of DDX3 within two minutes, and DDX3 inhibitors block the rapid NEU3 upregulation. The NEU3-TGF-β1-NEU3 positive feedback loop is activated within 5 minutes, and the remaining LAP fragment (after NEU3 activation of the latent TGF-β1 complex) appears to potentiate NEU3 upregulation. This feedback loop is blocked by NEU3 inhibitors. These findings suggest that NEU3 is a rapidly acting amplifier of TGF-β1 signaling, likely through TGF-β1-mediated dephosphorylation of DDX3, and this mechanism may be associated with an early tissue-healing response mechanism that drives the initiation and progression of fibrosis.

## Introduction

Fibrotic disorders, characterized by a progressive buildup of scar tissue in an internal organ, contribute to nearly 45% of deaths in the United States [1, 2]. Idiopathic pulmonary fibrosis (IPF), a lung-specific fibrotic disorder, has a median survival of only 4.5 years after diagnosis [3]. While current treatments can slow disease progression, they do not reverse or stop fibrosis, and the prognosis for patients remains poor [4]. A key driver of fibrosis is the extracellular cytokine transforming growth factor beta1 (TGF-β1) [5, 6]. TGF-β1 is initially synthesized as an inactive pre-pro-protein with a large N-terminal prodomain called latency-associated peptide (LAP) [7, 8]. After cleavage of the signal peptide, TGF-β1 is trafficked to the endoplasmic reticulum where two TGF-β1 pro-proteins form a disulfide-linked homodimer [9]. After trafficking to the Golgi, proteases cleave the LAP prodomain, LAP and TGF-β1 are secreted, and LAP remains noncovalently associated with TGF-β1, forming the latent TGF-β1 complex [10–12]. The latent TGF-β1 complex binds to the extracellular matrix via a covalent interaction with latent TGF-β-binding protein (LTBP) where LAP sequesters TGF-β1 until activation [9, 10]. Multiple mechanisms can release active TGF-β1 from LAP [13–17]. Like many other secreted proteins, LAP is glycosylated [18–20], and many glycosylated structures have sialic acids at the distal end of the polysaccharide chain which help regulate various cellular mechanisms [21–24]. Sialidases, also known as neuraminidases, remove the terminal sialic acid from glycoconjugates, and desialylation of LAP can also release active TGF-β1 [25, 26].

There are four known mammalian sialidases, neuraminidase 1 through 4 (NEU1-NEU4), each with distinct substrate specificities and subcellular localizations [25, 27]. Among them, NEU3 is found intracellularly in endosomes, on the extracellular side of the plasma membrane, and under some conditions it may also be released from the membrane to the extracellular environment [27–31]. NEU3 has been implicated in a range of pathological processes across organ systems including intestinal inflammation and colitis [32], liver fibrosis [33], and atherosclerosis [34]. In a mouse model of intestinal inflammation and colitis, NEU3 deficiency protects against disease onset [32]. For liver fibrosis, patients with nonalcoholic fatty liver disease (NAFLD) and steatohepatitis (NASH) have increased desialylation of serum proteins, and NEU3 levels are increased in NAFLD patients and mice with carbon tetrachloride (CCl_4_)-induced liver fibrosis [33]. Treatment with the NEU3 inhibitor 2-acetylpyridine (2-AP) in this mouse model reduces CCl_4_ induced weight loss, inflammation, and desialylation in the liver [33]. NEU1 and NEU3 can trigger the initial phase of atherosclerosis by desialylating low-density lipoproteins (LDL) [34]. Conversely, in certain forms of cardiac fibrosis, NEU3 may initially play a protective role by responding to chronic hypoxia and activating cardioprotective hypoxia-induced pathways [35, 36]. NEU3 knockout mice do not develop bleomycin-induced fibrosis [37], and NEU3 aspiration is sufficient to induce pulmonary fibrosis in mice [29], indicating that NEU3 is both necessary and sufficient for pulmonary fibrosis in mice. In a mouse bleomycin model of pulmonary fibrosis, inhibition of NEU3 with general sialidase inhibitors such as 2,3-dide-hydro-2-deoxy-N-acetyl-neuraminic acid (DANA) and oseltamivir, and more specific NEU3 inhibitors such as 2-acetyl pyridine (2-AP) and 4-Amino-1-methyl-2-piperidinecarboxylic acid (AMPCA), inhibits pulmonary inflammation, decreases active TGF-β1 levels in the lungs, and decreases pulmonary fibrosis [26, 29, 30, 37]. NEU3 can contribute to fibrosis through three key mechanisms: 1) NEU3 desialylates the endogenous serum antifibrotic protein serum amyloid P (SAP), which inhibits SAP activity [38, 39], 2) NEU3 upregulates the pro-inflammatory cytokine IL-6 in human peripheral blood mononuclear cells, which in turn upregulates NEU3, functioning in a positive feedback loop [26, 37, 39], and 3) NEU3 desialylates LAP, releasing TGF-β1 from latent TGF-β1, and the released active TGF-β1 upregulates NEU3 translation and decreases NEU3 degradation, functioning in a second positive feedback loop [30, 40].

For about two-thirds of the proteins responsive to TGF-β1, regulation occurs at the transcriptional level through the classic multi-step process of 1) influencing a transcription factor, 2) nuclear translocation, 3) gene transcription, 4) mRNA processing, 5) mRNA export from the nucleus, and 6) translation [41, 42]. This multistep process imposes a temporal delay such that increases in protein levels typically occur after several hours. For instance, in chondrocytes, 5 ng/ml of TGF-β1 increases Sox9 after 2 hours but not at 30 minutes or 1 hour [43]. In human lung fibroblasts, 2 ng/ml of TGF-β1 increases alpha smooth muscle actin (α-SMA) at 12 hours but not at 1, 3, or 6 hours [44]. In human kidney cells, 10 ng/ml of TGF-β1 upregulates vimentin and Snail1 at 72 hours but not at 24 hours [45]. In human bronchial epithelial cells, 10 ng/ml of TGF-β1 increases N-cadherin at 12 hours but not at 2 hours, and increases vimentin and Snail1 at 48 hours but not at 2, 12, or 24 hours [46]. In contrast, the remaining one-third of proteins are controlled at the level of translation where the protein levels may be changed without having to complete steps 1-5, potentially allowing a relatively quicker response [47, 48].

In human lung fibroblasts, there are at least 182 proteins including NEU3 whose levels are increased by TGF-β1 without a corresponding increase in their mRNA levels, but with a shift in these mRNAs from free and monosome fractions to polysomes, indicating translational regulation [49]. Of the 182 proteins, 180 (including NEU3) share a 20-nucleotide motif. This motif is necessary and sufficient for TGF-β1-induced translation of NEU3 [49]. DEAD-box helicase 3 (DDX3) is an ATP-dependent RNA helicase that is upregulated in the lungs of mice with bleomycin-induced pulmonary fibrosis and in fibrotic lesions of IPF patients [49]. In response to TGF-β1, DDX3 increases its binding to the 20-nucleotide common motif [49]. DDX3 inhibition with the DDX3 inhibitor RK-33 [50] reduces TGF-β1 upregulation of NEU3 in human lung fibroblasts, and in mice RK-33 injections reduce bleomycin-induced lung inflammation and fibrosis, and lung tissue levels of DDX-3, TGF-β1, and NEU3 [49]. This suggests that DDX3 may play a key role in the TGF-β1 mediated regulation of NEU3.

In this report, we show that NEU3 is rapidly upregulated by TGF-β1 via a translation-dependent mechanism with intracellular and extracellular levels rising within 5 minutes, identifying NEU3 as one of the key early effectors of TGF-β1 signaling. TGF-β1 induces dephosphorylation of DDX3 within 2 minutes and the rapid upregulation of NEU3 is blocked by pharmacological inhibition of DDX3 with RK-33. Additionally, we show the previously hypothesized NEU3-TGF-β1 positive feedback loop [26, 39, 40] is activated and amplified within 5 minutes, and that the LAP component of the activated latent TGF-β1 complex appears to contribute to the upregulation NEU3. This feedback loop is blocked by the NEU3 inhibitors 2-AP and AMPCA. This suggests a mechanism where TGF-β1 rapidly dephosphorylates DDX3, which increases the translation of *NEU3* mRNA which in turn rapidly activates TGF-β1 in a positive feedback loop to generate a very fast wound-healing response that when dysregulated in an active state can significantly contribute to fibrosis.

## Materials and Methods

### Primary human lung fibroblast cell culture

Eight human lung fibroblast (HLF) cell lines from 3 healthy males (M2, M3, NL-83), 1 healthy female (NL-44), 3 female IPF patients (IPF-36, IPF-8, F5), and 1 male IPF patient (IPF-32), were gifts from Dr. Carol Feghali-Bostwick, Medical University of South Carolina, Charleston, South Carolina, USA. Frozen cell lines were thawed and cultured in 6-well tissue culture dishes (10062-892, VWR, Radnor, PA) in DMEM (15-017-CV, Corning, Glendale, AZ) supplemented with 10% bovine calf serum (BCS) (SH30072.03, Cytiva, Wilmington, DE), 100 U/mL penicillin and 100 μg/mL -streptomycin (SV30010 Cytiva), and 2 mM glutamine (SH30034.01, Cytiva) (referred herein as “culture medium”), for 2-4 days at 37°C in a humidified 5% CO_2_ incubator, replacing the culture medium every 2 days. After reaching ∼60-80% confluence, the medium was removed, cells were rinsed twice with 1 ml of 1x phosphate buffered saline (PBS) (21-040-CV, Corning) before detaching with 0.5 ml of Accutase (AT-104, Innovative Cell Technologies, San Diego, CA). After detaching, Accutase was neutralized with 1 ml of culture medium, 14 ml of PBS was added, and cells were collected by centrifugation at 200 x g for 5 minutes. The cells were resuspended in culture medium to 1.5 x 10^4^ cells/ml, and added to 24-well plates (353047, Corning) at 500 µl/well or 96-well black μ-plates (89626, Ibidi Fitchburg, WI) at 200 µl/well. For all experiments, cells were used at passages 4–10, and no abnormal morphology differences were observed in this passage range. “Protein free medium” (DMEM/ 100 U/mL penicillin/ 100 μg/mL streptomycin/ 2 mM glutamine) was used for most experimental dilutions and as a negative control. Culture medium and protein free medium were warmed to 37°C before use.

### Coomassie staining of gels and western blot

HLF lines in 24-well plates were grown in culture medium for 2-4 days at 37°C in a humidified 5% CO_2_ incubator. After reaching ∼60-80% confluence, the liquid was removed, cells were washed twice with PBS, and 200 μl of protein free medium with or without 10 ng/ml of recombinant human active TGF-β1 (781802, BioLegend, San Diego, CA, diluted from the vendor 0.2 mg/ml stock in protein free medium) was added to the wells. At the indicated times, the liquid was removed, cells were washed once with 1x PBS, and cells were lysed *in situ* with 150 μL of Laemmli sample buffer with 10% 2-mercaptoethanol (6050 EMD Millipore Corporation, Norwood, OH) and left for 5 minutes. The liquid was pipetted up and down twice to collect all the cells and transferred to Eppendorf tubes. Samples were heated at 95°C for 5 minutes and 5 μl were loaded on two 4%-20% Tris-glycine Mini-Protean TGX gels (Bio-Rad, Hercules, CA), one for Coomassie staining and one for a western blot. The gels ran at 90 volts for 1.5 hours in 1x Tris/glycine/SDS (25mM, 192mM, 0.1% respectively) running buffer. For total protein evaluation of samples, one gel was stained with Coomassie staining solution (0.1% w/v Coomassie brilliant blue (0615-10G, Bio-Rad)/ 50% methanol/ 10% glacial acetic acid/ 40% water) for 1 hour with gentle shaking, and destained with destaining solution (10% glacial acetic acid, 40% ethanol, 50% water) overnight with gentle shaking (after the first hour, the gel was rinsed with water and the destaining solution was replaced). Coomassie-stained gels were imaged using a ChemiDoc XRS+ System (Bio-Rad) and the integrated density of lanes was measured using Image Lab software (Version 6.1, Bio-Rad).

For extracellular NEU3, following TGF-β1 exposure for five minutes, culture supernatants in 24-well plates were gently pipetted up and down, collected in Eppendorf tubes, and placed on ice. 50 μl of PBS was used to gently rinse cells and was collected and added to the supernatants. The conditioned medium was clarified by centrifugation at 200 x g for 10 minutes, the top 80% of the supernatant was collected and clarified at 10,000 x g for 10 minutes, and 20 μl was taken from the top of each tube, mixed with 4 μl of 6X Laemmli buffer, and processed as above.

For western blots, proteins were transferred to polyvinylidene difluoride membranes (88518, Thermo Fisher Scientific) in Tris/glycine/SDS buffer containing 20% methanol [30]. Membranes were blocked for 1 hour at room temperature with blocking buffer (2% IgG-free BSA (BSA-50, Rockland Immunochemicals) in Tris buffered saline with Tween 20 (TBST; 150 mM NaCl, 10 mM Tris-HCl pH 7.4 with 0.1% Tween 20 (0777-1L, VWR)). Membranes were incubated with either 1 μg/ml rabbit anti-NEU3 antibody (27879-1-AP, ProteinTech, Rosemont, IL) or 1 μg/ml rabbit anti-caldesmon-1 antibody (D5C80, Cell Signaling Technology, Danvers, MA), or where indicated, 1 μg/ml rabbit anti-DDX3 antibody (NBP2-67-121, Bio-Techne, Minneapolis, MN) diluted in blocking buffer overnight at 4°C on a platform rocker. The next day, blots were washed 3 times in TBST for 10 minutes each and incubated with peroxidase-conjugated donkey F(ab’)2 anti-rabbit secondary antibody (711-036-152, Jackson ImmunoResearch) at 400 ng/ml in blocking buffer for 1 hour. Blots were then washed 3 times in TBST for 10 minutes each and SuperSignal West Pico Chemiluminescence Substrate (34577, Thermo Fisher Scientific, Waltham, MA) was used following the manufacturer’s protocol to visualize the peroxidase using a ChemiDoc XRS+ System. The integrated density of bands was measured using FIJI (Version 1.54p) [51].

### Immunofluorescence and image analysis

HLF cell lines were added to 96-well black μ-plates at 2,000 cells per well and grown in culture medium for 24-48 hours at 37°C in a humidified 5% CO_2_ incubator. After reaching ∼50-60% confluence, the medium was removed, cells were washed twice with PBS, and treated with 200 μl of protein free medium with or without 10 ng/ml recombinant human active TGF-β1 at 37°C in a humidified 5% CO_2_ incubator. Other TGF-β1 concentrations or treatments were also used as indicated. Where indicated, recombinant human latent TGF-β1 (R&D systems, cat #299-LT/CF), was diluted in protein free medium and added to cells. Where indicated, a 50 kDa cutoff spin filter (UFC505024, MilliporeSigma, Burlington, MA) was pre-wet with 500 μl of culture medium, spun at 12,000 x g for 5 minutes, and the flow through and retentate were discarded. 500 μl of 100 ng/ml latent TGF-β1 was added to the spin filter and spun at 12,000 x g for 30 minutes. The retentate was reconstituted to 500 μl with protein free medium and diluted to the indicated concentrations. The NEU3 inhibitors 2-acetylpyridine (2-AP) (W325104-1KG-K, Sigma-Aldrich) [26, 30] and AMPCA (manufactured by Sundia MediTech Company Ltd) [26, 30] were made as 10 mM stocks in water, diluted in protein free medium, and added to cells simultaneously with TGF-β1. For staining, wells were washed with 200 μl of PBS and fixed with 100 μl of 4% paraformaldehyde (19210, Electron Microscopy Sciences, Hatfield, PA) in PBS for 10 minutes at room temperature. After fixing, the liquid was removed, cells were washed with 200 μl of PBS, and permeabilized with 100 μl of 0.5% Triton X-100 (J66624, Thermo Fisher) in PBS for 5 minutes. The liquid was removed, cells were washed with 200 μl of PBS, and then blocked with 200 μl of 1% IgG-free BSA in PBS (PBSB) for 1 hour at room temperature. After blocking, liquid was removed, and 100 μl of either rabbit anti-NEU3 antibody (OACA04338, Aviva Systems, San Diego, CA) or rabbit anti-caldesmon antibody (D5C80, Cell Signaling Technology,) was added at 1:1000 in PBSB and left overnight at 4°C. The next day. cells were washed 3 times for 5 minutes per wash with 0.05% Tween 20 in PBS (PBST). After washing, 100 μl of 1 μg/ml donkey F(ab′)_2_ anti-rabbit Alexa Fluor 488 secondary antibody (711-546-152, Jackson ImmunoResearch) in PBSB was clarified by centrifugation at 12,000 x g for 5 minutes and added for 30 minutes at room temperature. Cells were then washed 3 times for 5 minutes per wash with PBST. After washing, 50 μl of 2 μg/ml DAPI (Biolegend, cat #422801) in PBS was added to each well. Images were taken with an Eclipse Ti2 microscope (Nikon, Melville, NY) and analyzed with FIJI software. For each experiment, a control well stained with only secondary antibody was imaged to evaluate background fluorescence.

For each well, 2-5 images were taken, and 15-20 randomly selected cells were outlined with the freehand line tool on FIJI and the area and integrated density were measured. Integrated densities were then divided by the corresponding area and the intensity/ area of the cells stained with secondary antibody alone was subtracted. These 15-20 readings were then averaged. The average was then normalized to the experimental negative control (cells with no treatment, stained with anti-NEU3 or anti-caldesmon antibodies) as percent change, with the negative control as 100%.

### Transcription and translation inhibition and DDX3 inhibition with RK-33

Cycloheximide (94271, VWR), actinomycin D (BVT-0089-M005, Adipogen, San Diego, CA), or RK-33 (S8246, Selleck Chemicals, Houston, TX) were dissolved in DMSO to 10 mg/ml, 0.5 mg/ml, and 10 mM stocks respectively, and further diluted in protein free medium. HLF cell lines were seeded on 96-well black μ-Plates as above. After reaching ∼50-60% confluence, liquid was removed, cells were washed twice with PBS and pre-treated with 100 μl of either 50 μg/ml cycloheximide, 1 μg/ml actinomycin D, 10 μM RK-33, or an equal volume dilution of DMSO in protein free medium (control) at 37°C in a humidified 5% CO_2_ incubator. For actinomycin D and cycloheximide, after 10 minutes, liquid was removed, cells were washed with 200 μl of 1x PBS and treated with 200 μl of protein free medium with or without active TGF-β1 for indicated times, for RK-33, protein free medium with or without active TGF-β1 was added directly to the wells and left for 5 minutes. After indicated time points, wells were then washed with 1x PBS and stained by immunofluorescence as above.

### Phosphorylation of DDX3

HLF lines in 6-well plates were grown in culture medium for 4-5 days at 37°C in a humidified 5% CO_2_ incubator. After reaching ∼70-80% confluence, the liquid was removed, cells were washed twice with PBS, and 1 ml of protein free medium with or without 10 ng/ml of recombinant human active TGF-β1 was added to cells. At the indicated times, cells were washed once with PBS, detached with 0.5 ml of Accutase for 5 minutes at room temperature and collected in Eppendorf tubes. 1 mL of PBS was added to dilute the Accutase, and cells were collected by centrifugation at 200 x g for 5 minutes. Supernatants were removed and cells were resuspended and washed in 1.5 ml of PBS and collected by centrifugation at 200 x g for 5 minutes. The washes were repeated for a total of three washes with the last centrifugation at 500 x g. Supernatants were removed and phosphorylated proteins were isolated using TALON PMAC Phosphoprotein Enrichment Kit (635641, Takara Bio, San Jose, CA) following manufacturer’s protocol. Phosphorylated proteins and supernatants of total cell lysates were stained for DDX3 using western blots as described above.

### Identification of potential G-quadruplex regions in NEU3 mRNA

To identify possible G-quadruplex regions in the NEU3 mRNA, the NM_006656.6 NEU3 transcript from the National Center for Biotechnology Information (NCBI) database was analyzed using QGRS Mapper [52].

### Statistical analysis

Statistical analyses were performed using Prism 10 (GraphPad Software, La Jolla, CA, USA). Statistical significance between two groups was determined by unpaired t-test, or between multiple groups using ANOVA with Dunnett’s post-hoc test.

## Results

### TGF-β1 upregulates NEU3 levels within 2 minutes

We previously observed that a 48-hour exposure to 10 ng/ml of TGF-β1 in human lung fibroblasts causes some mRNAs (including *NEU3*) to shift from monosomes to polysomes and increase corresponding protein levels without an increase in mRNA levels, indicating translational regulation [49]. In contrast, in response to TGF-β1 other proteins (including caldesmon) have stable monosome-to-polysome ratios, and increased mRNA levels with a corresponding increase in protein level, indicating transcriptional regulation [49]. Translation regulation can result in a very rapid protein increase in response to a signal [53–55]. To determine how rapidly NEU3 is upregulated, we examined NEU3 expression as a function of time after exposure to TGF-β1, and compared this to caldesmon expression. Male and female human lung fibroblasts were treated with 10 ng/ml of TGF-β1, and NEU3 and caldesmon levels were evaluated using western blots normalized to Coomassie-stained gels of samples (**Figures 1A-D** and **Supplementary Figure 1**) and immunofluorescence (**Figures 1E-G**). As previously observed, due to two isoforms of NEU3, the anti-NEU3 antibodies detected two bands on western blots [26, 28, 31, 40] (**Figure 1A**). NEU3 protein levels increased at 2 minutes after TGF-β1 exposure on both western blots and immunofluorescence, whereas the earliest rise in caldesmon was seen at 4 hours (**Figures 1B, D and G**), similar to other TGF-β1 transcriptionally regulated proteins [43–46]. On the western blots, TGF-β1 caused NEU3 levels to increase to 183 ± 23% of baseline (100%) at 2 minutes (**Figure 1C**). Analysis of the data in supplemental data set 5 from Chen *et al.* [49] showed that after 48 hours of TGF-β1 exposure, the median increase of all proteins whose levels were increased by TGF-β1 was to 150% of the baseline level. The TGF-β1-induced increase of NEU3 at 2 minutes was thus higher than the median TGF-β1-induced protein increase at 48 hours. We observed that baseline levels of NEU3 decrease over time in protein free medium but not in serum-containing culture medium **(Supplementary Figures 2A and C-F)**, whereas caldesmon levels showed less of a decrease **(Supplementary Figures 2B, D, and G)**. This indicates that some serum factor maintains baseline NEU3 levels. At 24 hours in protein free medium, baseline levels of NEU3 decreased, but relative levels of NEU3 with TGF-β1 were increased compared to protein free medium control as reflected in the quantification of immunofluorescence and western blots (**Figures 1C, D, F, and G)**. These data indicate that compared to TGF-β1 upregulation of caldesmon, TGF-β1 can rapidly increase NEU3 levels.

**Figure 1.**
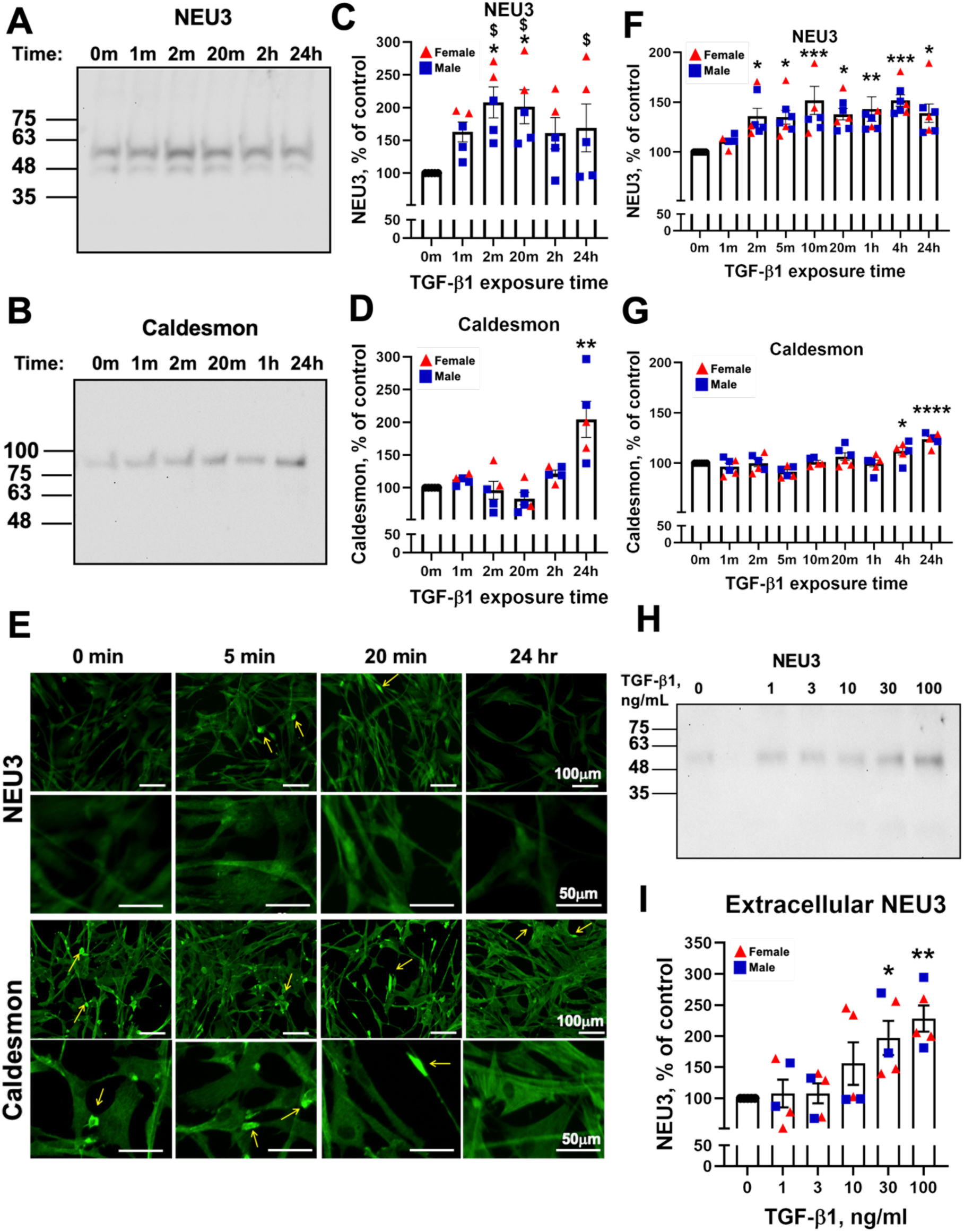
TGF-β1 upregulates intracellular and extracellular NEU3 levels within minutes. Human lung fibroblast cells were exposed to 10 ng/ml of TGF-β1 for the indicated times, lysed, and western blots were stained for **A)** NEU3 or **B)** caldesmon. m indicates minutes, h hours. The 0-minute samples were treated with protein free media only. Images are representative of 3 male and 2 female cell lines, each from a different donor. Molecular weights in kDA are indicated on the left. **C and D)** Quantification of western blots. Band densities were normalized to the integrated density of Coomassie-stained gels of total protein for each sample (**Supplementary Figure 1**). For NEU3, the 55 and 48 kDa bands were added together. For each cell line, 3 or more assays were done, and in each assay, after normalizing to Coomassie densities, values were normalized to the protein free media control at that time point (**Supplementary Figures 2A and B)**. The results were then averaged for each cell line. Blue squares indicate the averages for male cell lines and red triangles indicate the averages for female cells. Bars are mean ± SEM of the 5 averages (3 male and 2 female cell lines). * p<0.05, *** p<0.001 (One-way ANOVA, Dunnett’s test compared to the 0-minute control). $ p<0.05 (Unpaired t-test comparing male to female). **E)** Human lung fibroblasts were exposed to 10 ng/ml of TGF-β1 for the indicated times, fixed, and stained for NEU3 (rows 1 and 2) or caldesmon (rows 3 and 4) using immunofluorescence. The 0-minute samples were treated with protein free media only. Images are representative of 4 male and 3 female cell lines, each from a different donor. Rows 2 and 4 show a magnified view of images from rows 1 and 3 respectively. Bar is 100 µm for rows 1 and 3 and 50 µm for rows 2 and 4. Yellow arrows are examples of artifacts, detached cells, or abnormal appearing cells that were not included in fluorescence quantification. **F-G)** Quantification of immunofluorescence, m indicates minutes, h hours. Blue squares indicate male cell lines and red triangles indicate female cell lines. Values are normalized to protein free media control for each time point **(Supplementary Figures 2C-G).** Bars are mean ± SEM, n ≥ 7 (4 male and 3 female cell lines). * p<0.05 ** p<0.01, **** p<0.0001 (One-way ANOVA, Dunnett’s test compared to the 0-minute control). **H)** Human lung fibroblast cells were exposed to the indicated concentrations of TGF-β1 for 5 minutes. After pipetting up and down gently, liquid was collected off cells, and stained for NEU3 on western blots. Molecular weights in kDA are indicated on the left. **I)** Quantification of western blots. Band densities were normalized to the integrated density of Coomassie-stained gels of total protein for each sample, then values were normalized to the 0 ng/ml control. Blue squares indicate the averages for male cell lines and red triangles indicate the averages for female cells. Bars are mean ± SEM of the 5 averages (2 male and 3 female cell lines). * p<0.05, ** p<0.01 (One-way ANOVA, Dunnett’s test compared to the 0 ng/ml control).

### A 5-minute TGF-β1 exposure upregulates extracellular NEU3

NEU3 is found intracellularly in endosomes, on the extracellular side of the plasma membrane, and under some conditions it may also be released from the membrane to the extracellular environment [29–31]. To determine if a brief TGF-β1 exposure increases extracellular NEU3, human lung fibroblasts were exposed to TGF-β1, and western blots of the conditioned media were stained for NEU3. Only one band of NEU3 was observed in the cell culture supernatants (**Figure 1H**), which is consistent with previous observations [29, 31, 40]. Extracellular NEU3 protein levels increased at 5 minutes with 30 and 100 ng/ml of TGF-β1 (**Figures 1H** and **I**). These data suggest that in addition to rapidly increasing levels of intracellular NEU3, TGF-β1 also rapidly increases levels of extracellular NEU3.

### The rapid TGF-β1-mediated upregulation of NEU3 occurs in the presence of a transcription inhibitor but not in the presence of a translation inhibitor

TGF-β1 can upregulate NEU3 protein without a corresponding increase in *NEU3* mRNA levels [49]. To test the hypothesis that rapid NEU3 upregulation is independent of transcriptional control, we observed the NEU3 response to TGF-β1 in the presence of the transcription inhibitor actinomycin D (Act D) and the translation inhibitor cycloheximide (CHX) [51, 56]. Fibroblasts were pre-treated with either Act D or CHX, or a DMSO control, and then stimulated with 10 ng/ml of TGF-β1. When pre-treated with Act D, levels of the apparently transcriptionally regulated protein caldesmon did not significantly increase in response to TGF-β1 (**Figures 2A** and **B**). In contrast, TGF-β1 exposure rapidly upregulated NEU3 both in the Act D pre-treated and control conditions (**Figures 2A, C**). In cells treated with CHX [57], TGF-β1 did not significantly upregulate NEU3 (**Figure 2D**). In the DMSO control conditions, caldesmon levels increased sooner (20 minutes) than the previously observed increase at 4 hours (**Figure 1G**). This may be due to the presence of DMSO, which increases transcription through multiple mechanisms including weakening histone-histone interactions and relieving chromatin-mediated repression [58] as well as altering RNA polymerase conformation enhancing its initiation activity [59, 60].

**Figure 2.**
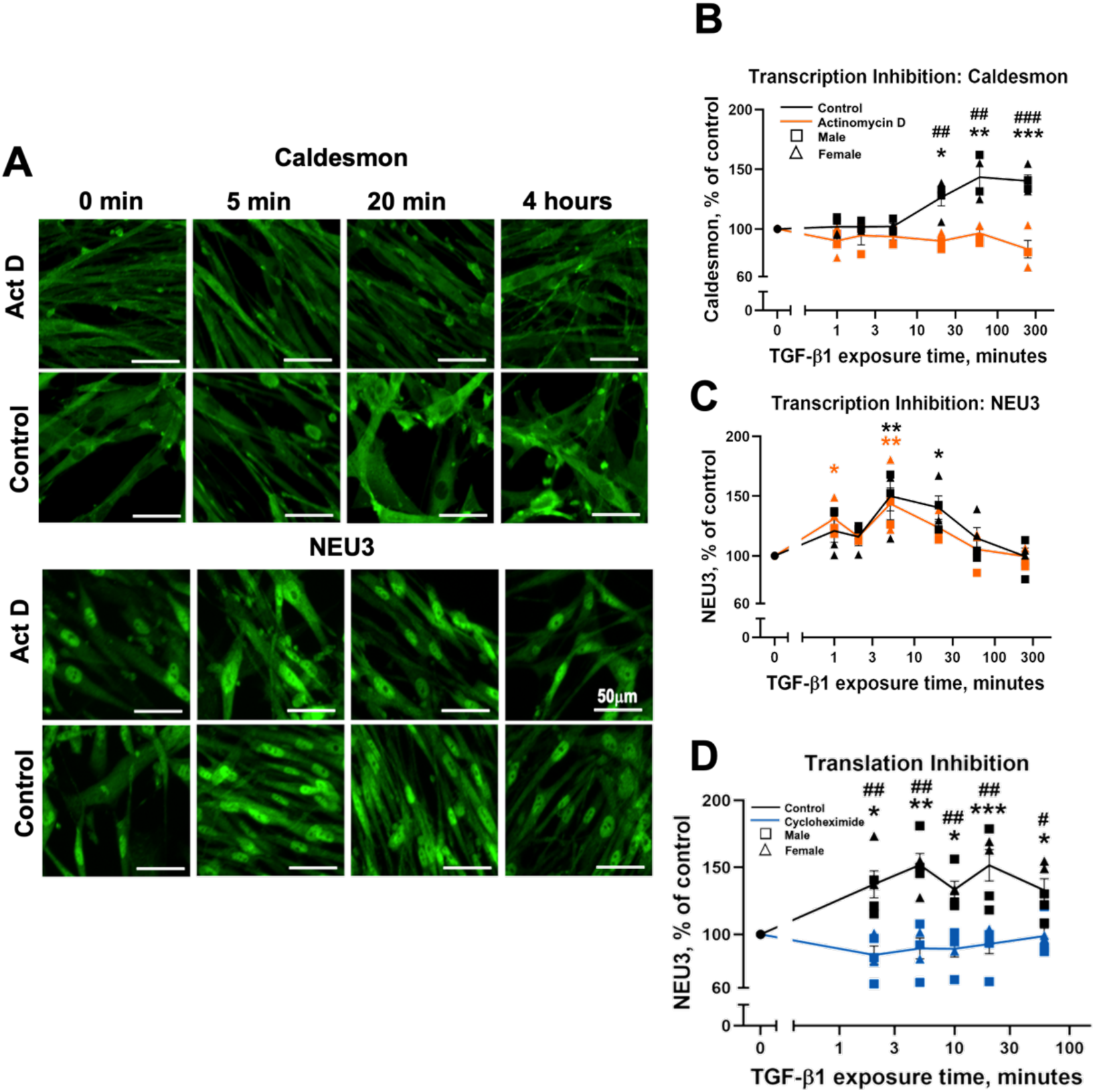
TGF-β1 upregulation of NEU3 occurs in the presence of a transcription inhibitor. **A)** Human lung fibroblasts were pre-incubated with either 1 µg/ml of the transcription inhibitor actinomycin D (Act D) or DMSO buffer control for 10 minutes. Cells were then washed and exposed to 10 ng/ml of TGF-β1 for the indicated times, fixed, and stained for caldesmon or NEU3 using immunofluorescence. The 0-minute samples were treated with protein free media only. Images are representative of 2 male and 2 female cell lines. Bars are 50 µm. **B and C)** Quantification of immunofluorescence. Actinomycin D treated cells are shown in orange and control cells are shown in black, squares indicate male cells and triangles indicate female cells. **D)** Human lung fibroblasts were pre-incubated with either 50 µg/ml of the translation inhibitor cycloheximide or buffer control for 10 minutes. Cells were then treated with TGF-β1 and NEU3 staining was quantified as in **A**. Cycloheximide treated cells are shown in blue and control treated cells are shown in black. Squares indicate male cells and triangles indicate female cells. Values are normalized to the 0-minute control. Bars are mean ± SEM, n=4 (2 male and 2 female cell lines). * p<0.05 ** p<0.01, *** p<0.001 (One-way ANOVA, Dunnett’s test compared to the 0-minute control). # p<0.05 ## p<0.01 ### p<0.001 (Unpaired t-test comparing actinomycin D or cycloheximide to control for each time point).

Notably, NEU3 control levels did not change with DMSO, supporting the idea that TGF-β1 upregulation of NEU3 is not mediated by a transcription mechanism. Overall, these data support the idea that TGF-β1 upregulation of NEU3 is mediated by translation and does not require TGF-β1 regulation of transcription.

### The DDX3 inhibitor RK-33 inhibits NEU3 upregulation, and TGF-β1 dephosphorylates DDX3

TGF-β1 increases the binding of the RNA helicase DDX3 to a common 20 nucleotide motif which is found in 180 human lung fibroblast mRNAs (including *NEU3*) whose levels are not significantly affected by TGF-β1, but whose translation is upregulated by TGF-β1 [49]. RK-33 is a DDX3 inhibitor that attenuates bleomycin-induced pulmonary fibrosis in young male mice [49]. To determine whether DDX3 inhibition attenuates the rapid upregulation of NEU3, male and female HLFs were pre-treated with RK-33, then stimulated with TGF-β1 for 5 minutes, and NEU3 was measured. As observed above, a 5-minute treatment with TGF-β1 increased NEU3 levels, and RK-33 blocked this upregulation (**Figure 3A**). Changes in phosphorylation state are rapid and often involved in signaling cascades [61]. To determine if TGF-β1 affects DDX3 phosphorylation, HLFs were treated with 10 ng/ml of TGF-β1. TGF-β1 dephosphorylated DDX3 within 2 minutes without significantly changing DDX3 levels (**Figures 3B-D and Supplementary Figure 3**). Together, these data suggest that the rapid NEU3 upregulation by TGF-β1 may be mediated by DDX3, and specifically by DDX3 dephosphorylation.

**Figure 3.**
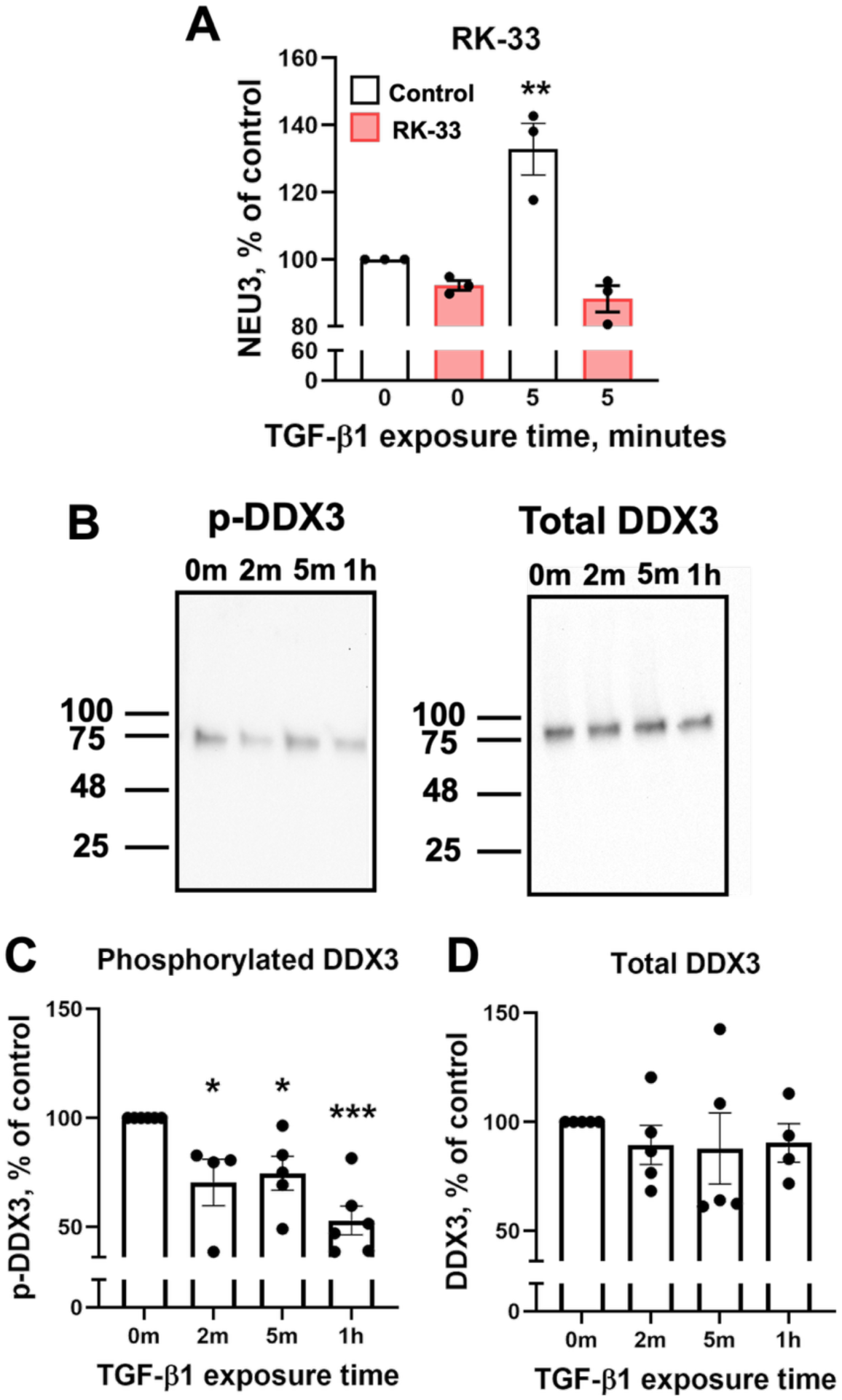
The DDX3 inhibitor RK-33 blocks the 5-minute NEU3 upregulation, and TGF-β1 induces rapid dephosphorylation of DDX3. **A)** Human lung fibroblast cells were pre-incubated with either 10 µM of RK-33 or an equal volume dilution of DMSO in protein free medium for 10 minutes. Protein free medium with or without TGF-β1 was added to wells (final TGF-β1 concentration 10 ng/ml) for 5 minutes. Cells were then washed, fixed, and stained for NEU3 using immunofluorescence. RK-33 treated cells are indicated with red bars. Values are normalized to 0-minute controls. Bars are mean ± SEM, n=3 (1 male and 2 female cells lines). ** p<0.01 (One-way ANOVA, Dunnett’s test compared to 0-minute control). **B**) Human lung fibroblasts were treated with 10 ng/ml of recombinant human active TGF-β1 for the indicated times. m indicates minutes, h hour. The 0-minute samples were treated with protein free media only. Phosphorylated proteins were isolated using TALON PMAC magnetic beads. Phosphorylated proteins and supernatants of total cell lysates were stained for DDX3 using western blots. Blots are representative of 5 donors (4 male and 1 female). **C and D)** Quantification of western blots. Band densities of phosphorylated and total DDX3 were normalized to the integrated density of silver- or Coomassie-stained gels respectively of total protein for each sample (**Supplementary Figure 3)**, then values were normalized to the 0-minute control. Bars are mean ± SEM, n=5 (4 male and 1 female). * p<0.05, *** p<0.001 (One-way ANOVA, Dunnett’s test compared to 0-minute control).

### NEU3 functions in a positive feedback loop with TGF-β1

In the latent TGF-β1 complex, the LAP protein is sialylated [18–20], and NEU3 desialylates LAP [26]. This desialylation causes the release of active TGF-β1 from LAP sequestration [26]. Based on these observations, we hypothesized that NEU3 functions in a positive feedback loop with TGF-β1 where NEU3 releases active TGF-β1 from the latent TGF-β1 complex, and the released TGF-β1 increases levels of NEU3 [26, 39, 40]. A prediction of this positive feedback amplification model is that for cells exposed to a low concentration of active TGF-β1, which causes a slight upregulation of NEU3, addition of latent TGF-β1 would allow the slightly increased levels of extracellular NEU3 to activate the latent TGF-β1, releasing active TGF-β1, causing further upregulation of NEU3. To test this hypothesis, we first examined the effect of active TGF-β1 on NEU3 upregulation and observed that at 1 ng/ml and below, although there was a slight upward trend, active TGF-β1 did not significantly increase NEU3 at 5 minutes as assessed by immunofluorescence (**Figure 4A**). Similarly, latent TGF-β1 alone (grey line in **Figure 4B**) upregulated NEU3 at 3, 10, and 30 ng/ml, but 1 ng/ml and lower concentrations of latent TGF-β1 did not significantly increase NEU3 (**Figure 4B**). Next, 0.1, 0.3, or 1 ng/ml of active TGF-β1 (which did not significantly upregulate NEU3) were mixed with a range of concentrations of latent TGF-β1 and added to fibroblasts for 5 minutes (**Figure 4B**). In the presence of latent TGF-β1, all three of these active TGF-β1 concentrations upregulated NEU3, with higher concentrations of active TGF-β1 needing less latent TGF-β1 for NEU3 upregulation, supporting the positive feedback model (**Figure 4B**). The commercial latent TGF-β1 preparation has a small amount of active TGF-β1 already present [62]. To test if the ability of latent TGF-β1 to upregulate NEU3 is due to this active TGF-β1 contaminant (∼22 kDa), the commercial latent TGF-β1 (∼75 kDa) was filtered using a 50 kDa cutoff spin filter and the retentate was added to human lung fibroblasts. Cells were treated with either filtered latent TGF-β1 alone or filtered latent TGF-β1 with 0.3 ng/ml of active TGF-β1. The filtered latent TGF-β1 upregulated NEU3 at 100 ng/ml (**Figure 4C**) but not at 3, 10, or 30 ng/ml as seen with the unfiltered commercial preparation (**Figure 4B**), suggesting that the NEU3 upregulation at lower concentrations of unfiltered latent TGF-β1 was due to the active TGF-β1 contaminant. Addition of 0.3 ng/ml of active TGF-β1 to 0.1 or 0.3 ng/ml of filtered latent TGF-β1 upregulated NEU3 (**Figure 4C**). As observed with other TGF-β1 mediated effects, high concentrations of active TGF-β1, unfiltered latent TGF-β1, or combinations did not significantly upregulate NEU3 [63, 64]. Although 3 ng/ml of active TGF-β1 alone is needed to upregulate NEU3 expression (**Figure 4A**), a combination of 1 ng/ml of active TGF-β1 with only 0.01 ng/ml of latent TGF-β1 was sufficient to significantly increase NEU3. This suggests that even if all of the active TGF-β1 were released from LAP in the latent TGF-β1 complex, the total concentration of active TGF-β1 would still remain below the 3 ng/ml threshold. Increased levels of NEU3 with a total possible TGF-β1 concentration below 3 ng/ml were also observed for the 0.1 and 0.3 ng/ml active TGF-β1 combinations (**Figures 4B** and **C**). This suggests the LAP component of the activated latent TGF-β1 complex may exert its own bioactivity to contribute to NEU3 upregulation that is independent of TGF-β1 (independent functions of LAP have been observed in other instances [13, 65–67]). These data support the idea that NEU3 activates latent TGF-β1 and this upregulates NEU3 in a positive feedback loop, and further suggest that the LAP remaining after NEU3 activation of latent TGF-β1 potentiates NEU3 upregulation.

**Figure 4.**
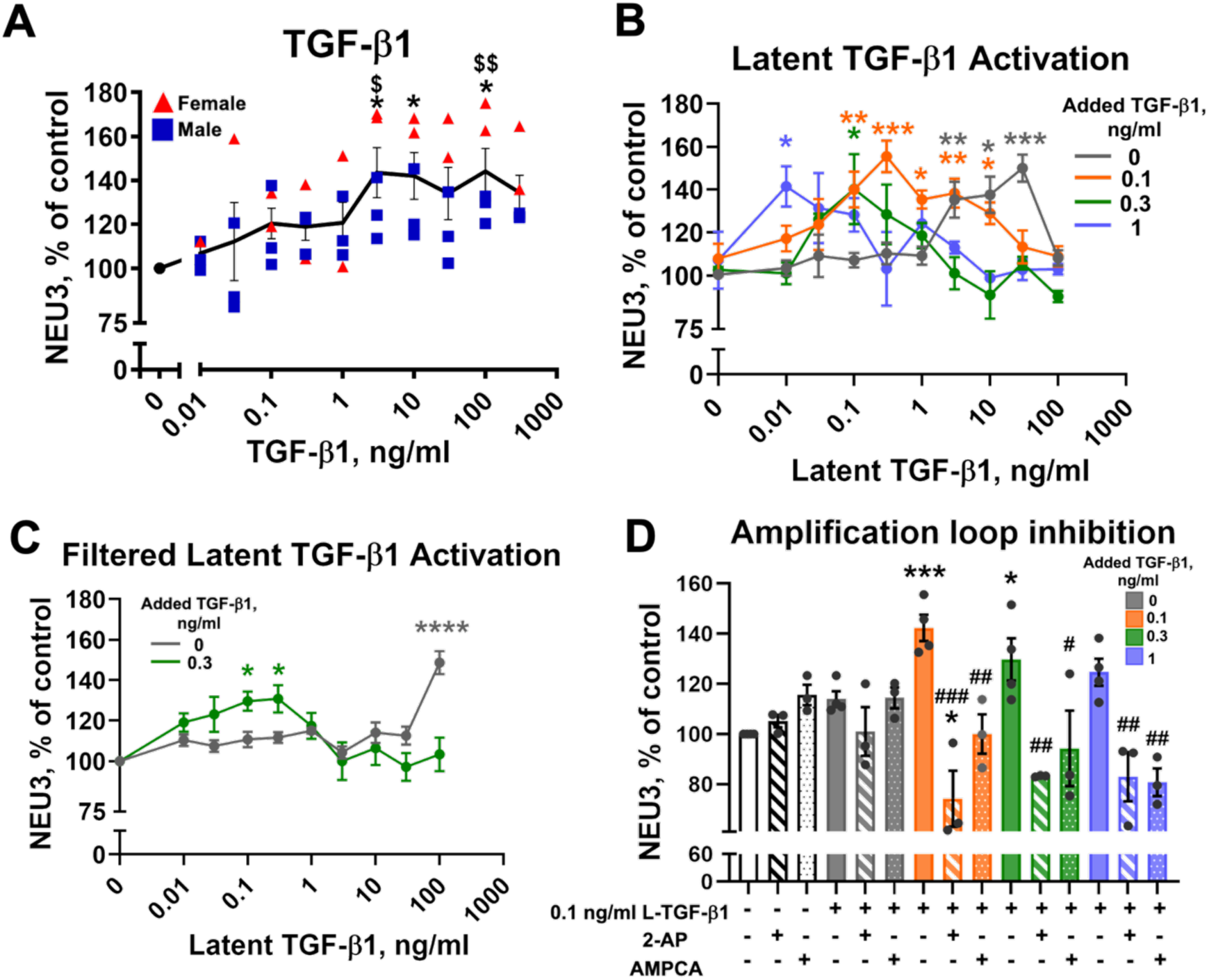
NEU3 functions in a positive feedback loop with TGF-β1 that is blocked by NEU3 inhibitors. **A)** Human lung fibroblast cells were exposed to the indicated concentrations of TGF-β1 for 5 minutes, fixed, and stained for NEU3 using immunofluorescence. Blue squares indicate male cells, and red triangles indicate female cells. Values are normalized to 0 ng/ml control. Bars are mean ± SEM, n=5 (3 male and 2 female cells lines). * p<0.05 (One-way ANOVA, Dunnet’s test compared to 0 ng/ml control). $ p<0.05, $$ p<0.01 (Unpaired t-test comparing male to female). **B)** Human lung fibroblast cells were exposed to 0, 0.1, 0.3 or 1 ng/ml of TGF-β1 (indicated with grey, orange, green, and blue respectively) and the indicated concentrations of latent TGF-β1 for 5 minutes, fixed, and stained for NEU3 using immunofluorescence. **C)** Human lung fibroblasts were exposed to 0 or 0.3 ng/ml of TGF-β1 (indicated with grey and green respectively) and the indicated concentrations of the retentate of latent TGF-β1 (purified with a 50 kDa spin filter) for 5 minutes, fixed, and stained with NEU3 using immunofluorescence. **D)** Human lung fibroblast cells were exposed to 0.1 ng/ml of latent TGF-β1 (L-TGF-β1) with either 0, 0.1, 0.3 or 1 ng/ml of TGF-β1 (indicated with grey, orange, green, and blue fill respectively, white fill indicates no latent TGF-β1 or added TGF-β1) in the presence or absence of NEU3 inhibitors: 2-AP and AMPCA (indicated with stripes and dots respectively). Cells were fixed and stained for NEU3 using immunofluorescence. For **B**-**D**, values are normalized to media control. For **B** and **D** bars are mean ± SEM, n=3 (2 male and 1 female cell line) for **C** bars are mean ± SEM, n=4 (2 male and 2 female cell lines). * p<0.05, ** p<0.01, *** p<0.001 **** <0.001 (One-way ANOVA, Dunnet’s test compared to media control). # p<0.05, ## p<0.01, ### p<0.001 (Unpaired t-test comparing with and without inhibitor for each concentration of TGF-β1).

### The NEU3-TGF-β1 feedback loop is inhibited by the NEU3 inhibitors 2-AP and AMPCA

The human and mouse NEU3 inhibitors 2-AP and AMPCA inhibit pulmonary inflammation, decrease levels of active TGF-β1 in the lungs, and decrease pulmonary fibrosis in a mouse bleomycin model [26]. To test if the NEU3 inhibitors influence the latent TGF-β1 amplification effect we observed, the latent TGF-β1 amplification experiment was repeated in the presence and absence of 2-AP and AMPCA. 0.1 ng/ml of latent TGF-β1 was mixed with the three suboptimal concentrations of active TGF-β1 (0.1, 0.3, 1 ng/ml) in the presence and absence of 10 nM 2-AP or AMPCA for 5 minutes and NEU3 levels were measured by immunofluorescence. In protein free media alone and 0.1 ng/ml of latent TGF-β1 alone, 2-AP and AMPCA did not significantly change the levels of NEU3 (**Figure 4D**). In contrast, both 2-AP and AMPCA blocked the amplification effect of NEU3 for all three active TGF-β1 concentrations (0.1 ng/ml latent TGF-β1 + 0.1, 0.3, and 1 ng/ml active TGF-β1). For unknown reasons, 0.1 ng/ml active TGF-β1 + 0.1 ng/ml latent TGF-β1 + 2-AP decreased NEU3 levels below baseline (**Figure 4D**). These data indicate that the NEU3 inhibitors 2-AP and AMPCA can block the NEU3-TGF-β1 feedback loop, inhibiting the upregulation of NEU3.

## Discussion

In this report, we show that unlike transcriptionally regulated proteins that increase after several hours of TGF-β1 exposure, NEU3 protein levels rise rapidly within 5 minutes both intracellularly and extracellularly. The rapid upregulation of NEU3 is independent of new transcription, supporting the idea that TGF-β1-induced upregulation of NEU3 is regulated on the translational level. In line with previous studies demonstrating a role of DDX3 in NEU3 translational regulation [49], we find that the rapid NEU3 upregulation is blocked by inhibiting DDX3 with RK-33. TGF-β1 induces DDX3 dephosphorylation within 2 minutes, suggesting that DDX3 dephosphorylation may be associated with the rapid NEU3 upregulation (**Figure 5**). For unknown reasons, we observed that in some cases, female cell lines had a slightly stronger NEU3 response than male cell lines following TGF-β1 stimulation. NEU3 activates latent TGF-β1, and the released active TGF-β1 upregulates NEU3 in a rapid positive feedback loop. This upregulation is potentiated by the LAP peptide after NEU3 activates latent TGF-β1. This loop can be inhibited by the NEU3 inhibitors 2-AP and AMPCA, supporting the direct role of NEU3 in this loop (**Figure 5**).

**Figure 5.**
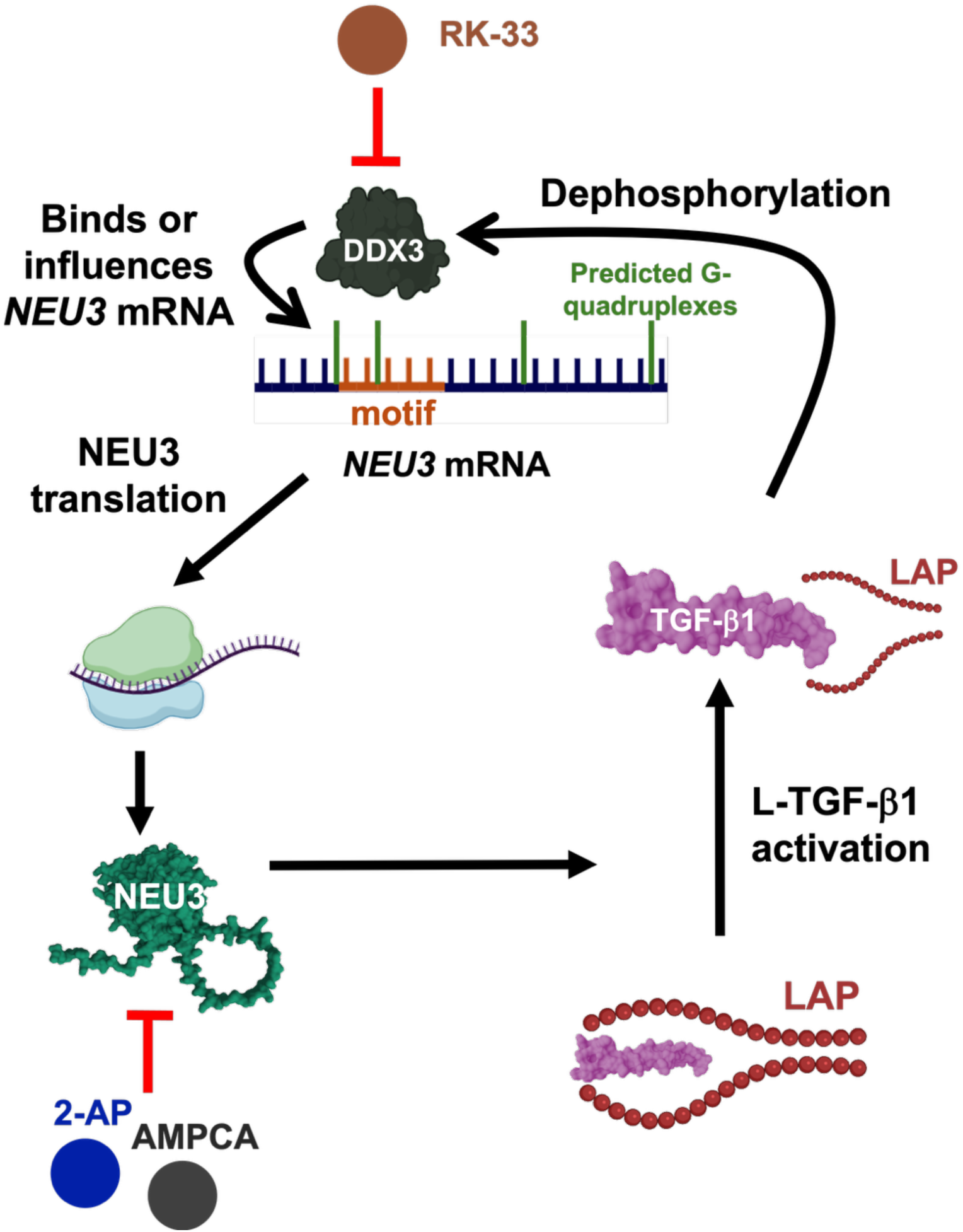
Proposed model of rapid NEU3 upregulation. Active TGF-β1 dephosphorylates DDX3 and increases the interaction of DDX3 with *NEU3* mRNA either by increasing binding to the common motif in *NEU3* mRNA, found in 179 other proteins that are translationally regulated by TGF-β1 (orange), or to G-quadruplex structures (green; the image labels 4 of the 32 predicted G-quadruplex structures). DDX3 stimulates rapid *NEU3* translation. Once translated and released from cells, NEU3 activates latent TGF-β1 (L-TGF-β1), releasing active TGF-β1 from its complex with LAP. TGF-β1, potentiated by LAP, upregulates NEU3 translation leading to a positive feedback loop. DDX3 is inhibited by RK-33, and NEU3 is inhibited by 2-AP and AMPCA, all three of which can disrupt this loop and the rapid 5-minute upregulation of NEU3. This figure was created using BioRender elements (https://biorender.com/); structures of NEU3 and TGF-β1 were predicted using AlphaFold 3 [68].

Fibrosis appears to be due to constitutively activated wound-healing mechanisms [48]. From an evolutionary standpoint, a rapid wound-healing response is crucial to minimize infection and further damage [6]. A small amount of positive feedback can significantly increase the sensitivity and response time of a system [69]. TGF-β1 is pre-positioned in the latent form in the extracellular environment, ready for rapid activation [9, 10]. In incisional wounds, active TGF-β1 is detectable within minutes [70], and subsequent TGF-β1-mediated changes in phosphorylation of downstream effectors are also detected within minutes [71, 72]. Together, this supports the idea that the rapid TGF-β1-NEU3-TGF-β1 positive feedback loop mechanism evolved to contribute to a rapid wound-healing response.

For translation to be rapidly upregulated without de novo *NEU3* mRNA synthesis, the *NEU3* transcript is likely pre-made and stored in the cytoplasm halted at the level of translation, as seen with other RNAs [73, 74]. For instance, in yeast, in preparation for stressful conditions, basal levels of mRNAs are transcribed and stored in the cytoplast. Under stressful conditions, these mRNAs are translated within minutes. This is seen for the yeast proteins GCN4 [75–77], HAC1 [78], and ICY2 [79, 80]. Similar mechanisms can be seen in human cells, with mRNAs that are stored in stress granules [81]. Together with our current findings, these results highlight a potential mechanism where *NEU3* mRNA is pre-made and stored in the cytoplasm, halted at the level of translation. Upon activation of latent TGF-β1, the released active TGF-β1 and LAP rapidly induce a translational regulator (such as DDX3) which activates the translation of *NEU3* mRNA within 5 minutes to form the TGF-β1-NEU3 part of the TGF-β1-NEU3-TGF-β1 positive feedback loop. NEU3 then exerts its enzymatic activity to activate TGF-β1 that is pre-positioned at the extracellular matrix, generating the NEU3-TGF-β1 part of the positive feedback loop for a rapid wound healing response. After release of the active TGF-β1, the LAP component may exert independent bioactivity to rapidly upregulate NEU3, and since LAP desialylation can release active TGF-β1 [26], it is possible that the desialylated form acquires a distinct signaling property that acts at an unknown receptor to reinforce the TGF-β1-NEU3-TGF-β1 feedback loop.

Changes in phosphorylation are one of the most rapid and effective regulatory mechanisms in signaling cascades [82, 83]. Our data suggests that TGF-β1 may induce dephosphorylation of DDX3, which in turn upregulates NEU3. Since dephosphorylation is a first-order reaction, depending only on the concentration of the phosphorylated substrate, it can proceed more rapidly than phosphorylation, which is a second-order reaction requiring both the substrate and ATP [84]. This kinetic difference often results in faster signal termination or activation through phosphatase activity. Similar rapid dephosphorylation events have been observed in other pathways such as MAPK signaling cascades, where phosphatase-mediated responses can occur within seconds [82].

Our previous work demonstrated that TGF-β1 enhances DDX3 binding to the common 20-nucleotide motif (shared among several mRNAs regulated by TGF-β1 on the translational level) after 1 hour of TGF-β1 stimulation but not at 30 minutes [49]. Although DDX3 binding to the motif showed an upward trend at 30 minutes, this association was not statistically significant. The current finding that TGF-β1 induces DDX3 dephosphorylation within 2 minutes, and that RK-33 blocks the rapid NEU3 upregulation implies that DDX3 activity is essential for this process, and suggests that DDX3 may cause a rapid increase in NEU3 levels through a mechanism that does not involve binding to the 20-nucleotide motif. One possibility is that dephosphorylated DDX3 does bind the common 20-nucleotide motif at early time points, which was not previously detected. Alternatively, DDX3 may interact with a different region of the NEU3 transcript. DDX3 has multiple binding sites for RNA [85, 86], and preferentially binds structured RNAs with G-quadruplex elements through its intrinsically disordered N-terminal region [87]. The NEU3 transcript contains multiple predicted G-quadruplex structures (**Figure 5**), raising the possibility that DDX3 binds these regions to increase translation at early time points. Another possibility is that DDX3 dephosphorylation alters its interaction network or helicase activity in a way that indirectly enhances NEU3 translation without direct RNA binding, such as modulating components of the translation machinery. Together these models support a mechanism where rapid TGF-β1 dephosphorylation of DDX3 upregulates the translation of NEU3.

There are at least two reasons that the rapid positive feedback loop does not invariably turn on in response to a single extracellular molecule of NEU3 or TGF-β1. First, both NEU3 and TGF-β1 have finite lifetimes allowing, as with other extracellular factors, a basal equilibrium to be reached between release rate and degradation/uptake rate. Second, the TGF-β1 signal transduction pathway has negative feedback components such as Smad7 [88], developmental endothelial locus-1 (Del-1) [89], and the RelA subunit of nuclear factor κB (NF-κB) [90], causing a threshold effect where low levels of TGF-β1 essentially do not activate the pathway. We hypothesize that in a wound or in fibrosis, a sufficiently high level of the various factors that increase NEU3 levels activate the loop. Under normal conditions, a small amount of TGF-β1 causes a small upregulation of NEU3 which in turn increases the amount of TGF-β1 to a new steady state level where the efficiency of the system is controlled (loop gain less than 1). In this normal environment, a decreased level of the injury signals and/or upregulation of inhibitory signals indicate a completion of healing and cause the loop to return to quiescent levels of NEU3 and TGF-β1. In fibrosis, three possibilities that could cause the loop to stay in an activated state include an excess of signals that activate the loop (due to increased amount of signal or decreased degradation), insufficient levels of signals that inhibit the loop, or an increase in the efficiency of the loop where NEU3 more efficiently upregulates TGF-β1 and/or TGF-β1 more efficiently upregulates NEU3 (loop gain equal to or more than 1).

High concentrations of TGF-β1 do not upregulate NEU3, which likely reflects a tightly controlled protective mechanism, as seen in multiple biological systems. For example, hormones often operate within complex dose-dependent negative feedback loops where high levels of a downstream effector downregulates its own production or secretion [91]. Additionally, continuous receptor stimulation can trigger receptor downregulation through decreased affinity and internalization [92]. However, these mechanisms typically take longer than a few minutes. A rapid downregulation mechanism could include NEU3 degradation in response to high activator concentrations [93, 94], rapid phosphorylation of downstream effectors that could inhibit NEU3 translation [71, 72], analogous to short-term desensitization of G-protein coupled receptors where phosphorylation of the receptor by GPCR kinase recruits β-arrestin to rapidly shut off the response [95, 96], or rapid receptor phosphorylation and uncoupling which is seen with serine threonine receptors [97]. Although the mechanism remains unclear, this non-linear dose effect indicates that NEU3 upregulation is likely a controlled process with mechanisms in place to avoid overstimulation.

Together, this report supports a model where *NEU3* mRNA is stored in the cytoplasm, ready for translational activation for a very rapid wound healing response which is amplified by a positive feedback loop that may be stuck in an on state in fibrosis.

## Declaration of interests

RHG is an inventor on patent application for the use of DDX3 inhibitors and NEU3 inhibitors as therapeutics for fibrosis.

## Acknowledgements

We thank Darrell Pilling for helpful comments. Funding for this work came from the Texas A&M University’s start-up fund for Richard Gomer.

