## Supplementary figures 1-3 for "Neuraminidase 3 acts in a rapid translation-based positive feedback loop to activate TGF-β1"

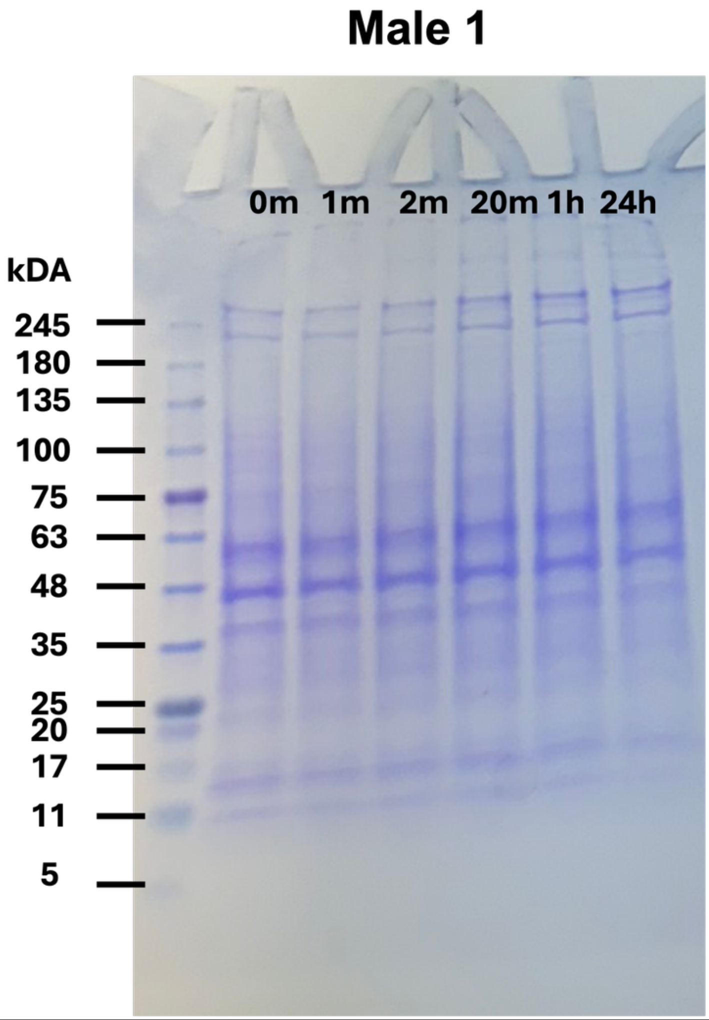


**Supplementary Figure 1. Coomassie-stained gel of total protein**. Human lung fibroblast cells were exposed to 10 ng/ml of TGF-β1 for the indicated times, lysed, and ran on gels stained with Coomassie to evaluate total protein, which was used to normalize the western blot bands in **Figure 1**. m indicates minutes, h hours. Image is representative of 3 male and 2 female cell lines used in **Figure 1**.

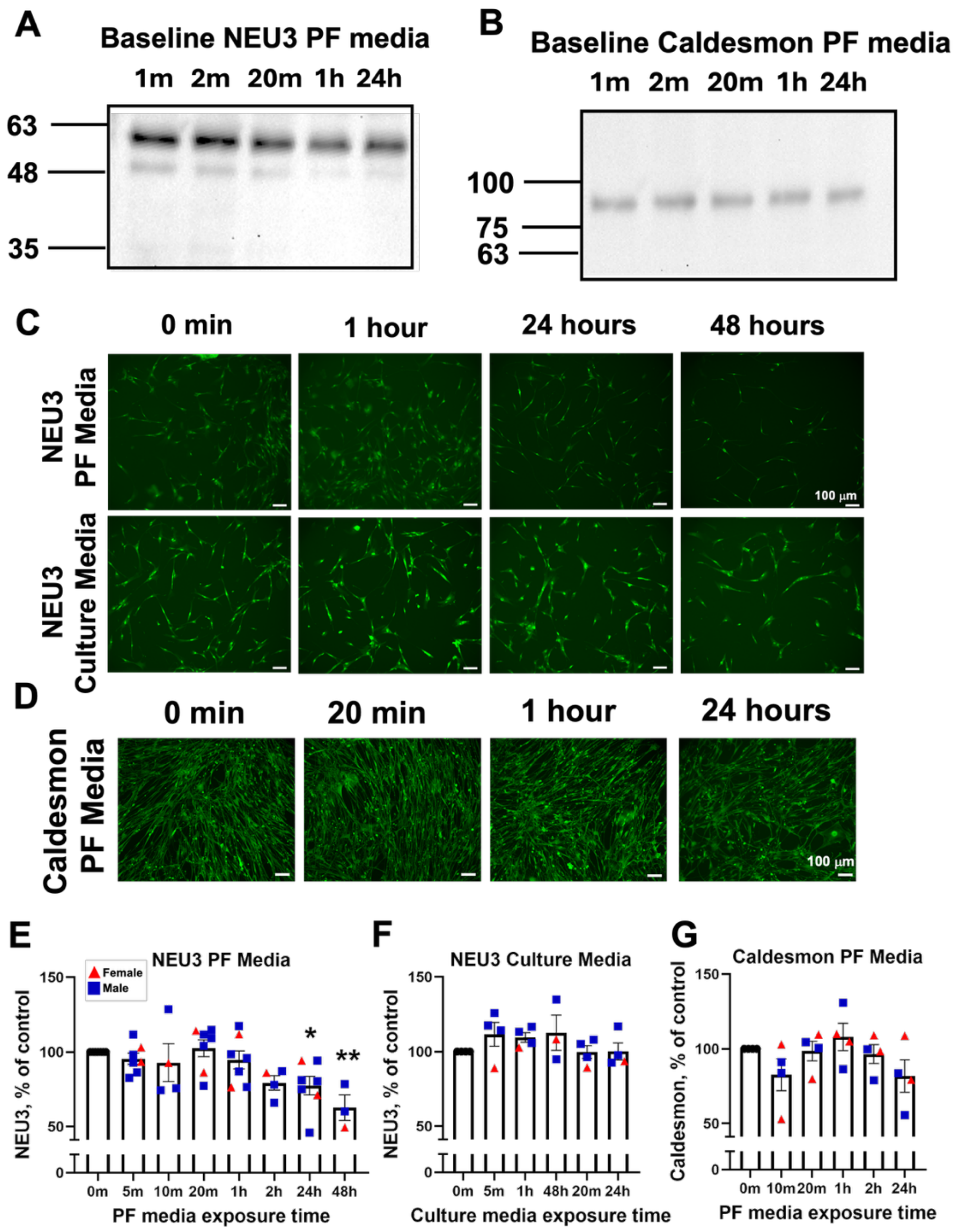


**Supplementary Figure 2. Baseline levels of NEU3 and caldesmon over time.** Human lung fibroblasts were incubated in protein free (PF) medium for the indicated times, lysed, and stained for **A)** NEU3 or **B)** caldesmon. m indicates minutes, h hours. Blots are representative of 6 donors (4 male and 2 female). Human lung fibroblasts were incubated in protein free (PF) medium (no serum) or culture medium (containing serum) for the indicated times, fixed, and stained for **C)** NEU3 or **D)** caldesmon by immunofluorescence. Bars are 100 µm. Images are representative of 7 donors (5 male and 2 female). **E)** Quantification of NEU3 in PF medium over time. **F)** Quantification of NEU3 in culture medium over time **G)** Quantification of caldesmon in PF medium over time. Values were normalized to the 0-minute control. Bars are mean ± SEM of the 4-7 cell lines. * $p<0.05$, ** $p<0.01$ (One-way ANOVA, Dunnett's test compared to the 0-minute control).

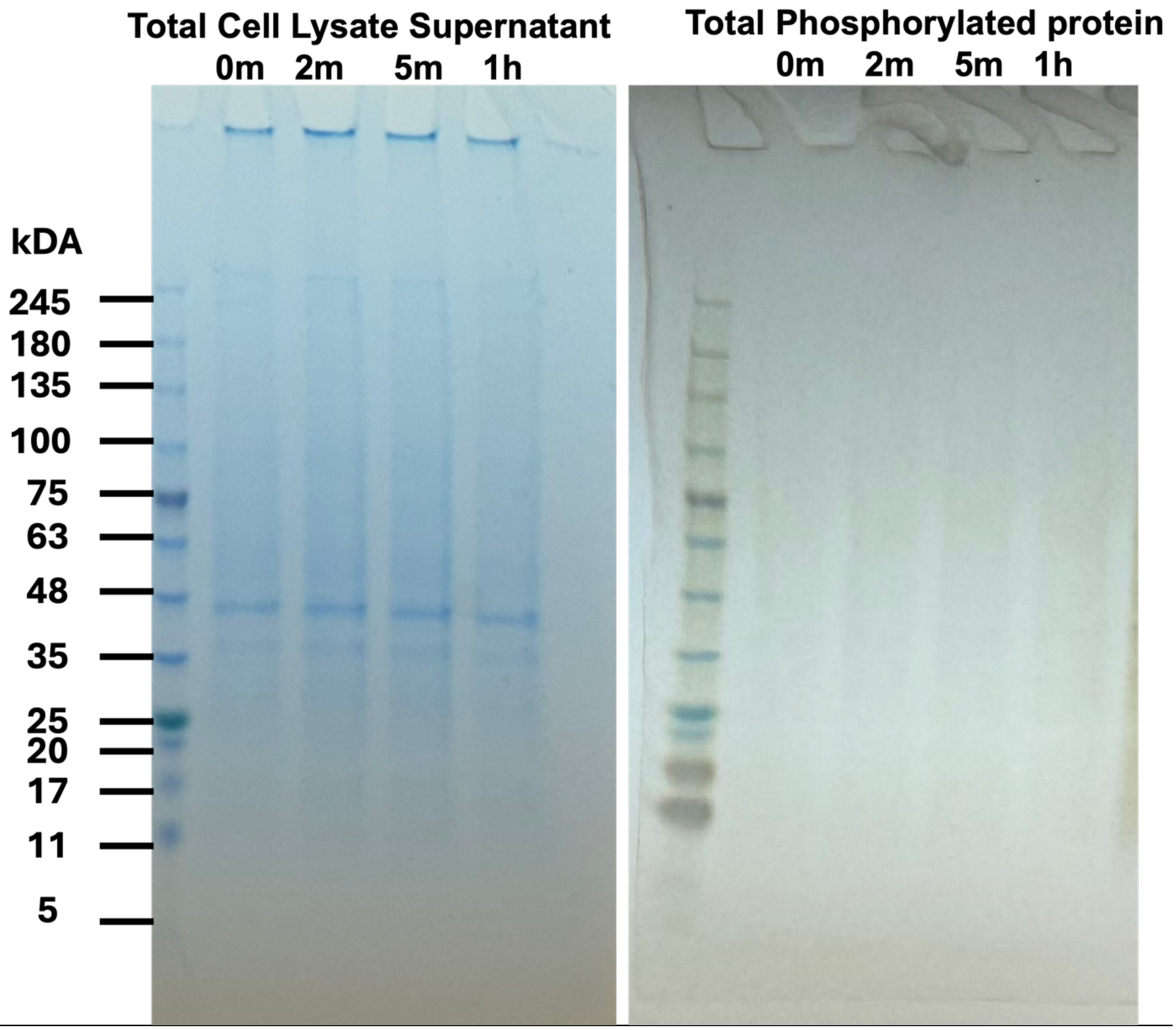


**Supplementary Figure 3. Coomassie- and silver-stained gels for total cell lysate and phosphorylated proteins**.

Human lung fibroblasts were treated with 10 ng/ml of recombinant human active TGF-β1 for the indicated times. The 0-minute samples were treated with protein free media only. Phosphorylated proteins were isolated using TALON PMAC magnetic beads. Total cell lysates (left) and total phosphorylated proteins (right) were run on gels stained with Coomassie and silver respectively. m indicates minutes, h hours. Image is representative of five cell lines used to normalize graphs in **Figure 3**.
